# The Pace of Modern Life, Revisited

**DOI:** 10.1101/2021.07.30.454364

**Authors:** Sarah Sanderson, Marc-Olivier Beausoleil, Rose E. O’Dea, Zachary T. Wood, Cristian Correa, Victor Frankel, Lucas D. Gorné, Grant E. Haines, Michael T. Kinnison, Krista B. Oke, Fanie Pelletier, Felipe Pérez-Jvostov, Winer Daniel Reyes-Corral, Yanny Ritchot, Freedom Sorbara, Kiyoko M. Gotanda, Andrew P. Hendry

## Abstract

Wild populations must continuously adapt to environmental changes or they risk extinction. Such adaptations can be measured as phenotypic rates of change and can allow us to predict patterns of contemporary evolutionary change. About two decades ago, a dataset of phenotypic rates of change in wild populations was compiled. Since then, researchers have used (and expanded) this dataset to look at microevolutionary processes in relation to specific types of human disturbances. Here, we have updated the dataset adding 5675 estimates of phenotypic changes and used it to revisit established patterns of contemporary evolutionary change. Using this newer version, containing 7338 estimates of phenotypic changes, we revisit the conclusions of four published articles. We then synthesize the expanded dataset to compare rates of change across different types of human disturbance. Analyses of this expanded dataset suggests that: I. a small absolute difference in rates of change exists between human disturbed and natural populations, II. harvesting by humans results in larger rates of change than other types of disturbances, III. introduced populations have increased rates of change, and IV. body size does not increase through time. Overall, findings from earlier analyses have largely held-up in analyses of our new dataset that encompass a much larger breadth of species, traits, and human disturbances. Lastly, we found that types of human disturbances affect rates of phenotypic change, and we call for this database to serve as a steppingstone for further analyses to understand patterns of contemporary evolution.

## INTRODUCTION

The rate at which populations can respond adaptively to environmental change will determine their ability to persist, thrive, and expand in a changing world. These adaptative responses are determined by changes in organismal phenotypes (as opposed to genotypes) – because phenotypes interface with the environment and thus are the direct determinants of fitness (Alberti et al., 2017; Endler, 1986; Hendry, 2017; Schluter, 2000). Throughout this paper we therefore use a definition of adaptive response that includes genetic and plastic contributions to phenotypic change, and, in many cases, we cannot differentiate between the two. Historically, such adaptive phenotypic changes were thought to be very slow, as exemplified by Charles Darwin’s statement that “*we see nothing of these slow changes in progress until the hand of time has marked the long lapse of ages*” (Darwin, 1859). This assumption began to crumble with the accumulation of studies documenting so-called “rapid evolution.” Some famous early examples of this phenomenon included industrial melanism in peppered moths (Kettlewell, 1973), body size in mice colonizing islands (Berry, 1964), body size and colour in house sparrows invading North America (Johnston & Selander, 1964), and resistance of plants to pollutants found in mine tailings (Antonovics & Bradshaw, 1970). In most of these early examples from nature, it was unclear whether the observed phenotypic changes were genetic as opposed to plastic. It was therefore game-changing when a series of common-garden experiments confirmed that at least some rapid phenotypic changes seen in nature are, in fact, genetically based (Al-Hiyaly, Mcneilly, & Bradshaw, 1990; Reznick, 1982; Stearns, 1983; Wu & Kruckeberg, 1985).

At the end of the 20^th^ century, it remained unclear if the documented cases of rapid phenotypic change were rare exceptions or the tip of the iceberg. Resolving this uncertainty required broader literature surveys and a quantitative standard for calculating and comparing rates of phenotypic change. A precedent already existed in the literature because paleontologists had long been calculating rates of phenotypic change in fossil time series (Gingerich, 1983, 1993; Haldane, 1949). Following that precedent, Hendry and Kinnison (1999) combed the literature for examples of phenotypic change on contemporary time scales and calculated rates of change using two classic metrics (“darwins” and “haldanes”). The authors concluded that “[…] *evolution as hitherto considered “rapid” may often be the norm and not the exception*” (Hendry & Kinnison, 1999). They further advocated use of the general term “contemporary evolution” because “rapid evolution” requires formal confirmation of exceptionally rapid rates. This original paper, and the follow-up analysis of Kinnison and Hendry (2001), opened the flood-gates to a series of influential papers analyzing the dataset of rates of phenotypic change to answer a series of evolutionary questions (Alberti et al., 2017; Crispo et al., 2010; Gorné & Díaz, 2019; Gotanda, Correa, Turcotte, Rolshausen, & Hendry, 2015; Hendry, Farrugia, & Kinnison, 2008; Hendry & Kinnison, 2001; Palkovacs, Wasserman, & Kinnison, 2011; Uyeda, Hansen, Arnold, & Pienaar, 2011; Westley, 2011). The studies also made the case that evolution operates on scales large enough to have ecological outcomes, leading the way into the field of eco-evolutionary dynamics (Des Roches et al., 2018; Fitzpatrick et al., 2015).

The number of studies available for calculating rates of phenotypic change has increased dramatically over the last decade or so. Compared to the last published version of the dataset that contained only populations in the wild (Alberti et al., 2017), the new dataset has increased from 1663 to 7338 rates, 89 to 214 studies, 175 to 1654 systems (Box 1), and 155 to 329 species. Hence, our first goal in the current paper is to present a new quality-controlled and much-expanded database of contemporary phenotypic changes in nature – The Phenotypic Rates of Change Evolutionary and Ecological Database (PROCEED) Version 5.0. This new database is publicly available at Dryad and https://proceeddatabase.weebly.com/. Our second goal was to use the new dataset to revisit and replicate previous analyses and conclusions based on earlier versions of the dataset. Specifically, we want to know if the effect sizes obtained in previous analyses have changed with the addition of new data. We ask four questions: I. Does the evidence still support the conclusion from Hendry et al., (2008) that phenotypic change is greater in human-disturbed systems than in more “natural” systems (see also Alberti et al., 2017)? II. Does the evidence still support the conclusion from Darimont et al., (2009) that phenotypic changes are most rapid when humans act as predators, such as during harvesting (see also Sharpe & Hendry 2009)? III. Does the evidence still support the conclusion from Westley (2011) that introduced populations do not drive particularly rapid phenotypic changes, relative to non-introduced populations? IV. Does the evidence still support the conclusion from Gotanda et al., (2015) that no evidence exists for microevolutionary trends toward increasing body size – as had been suggested to follow from “Cope’s Rule” (J. Kingsolver & Pfennig, 2004)?

The inclusion of 5675 additional data entries might change earlier conclusions for either of two reasons: biases in earlier data compilations such as underrepresented or missing taxa and disturbances, or statistical limitations including small sample sizes associated to taxonomic levels or types of studies (i.e., genetic; Box 1). To assess the first possibility, we re-analyze the new dataset using the same methods as the original authors: that is, Hendry et al., (2008), Darimont et al., (2009), Westley (2011), and Gotanda et al., (2015). Through this approach, we can assess if previous approaches yield similar conclusions following the accumulation of more data. To assess the second possibility, we use updated statistical models in a comprehensive approach to ask: V. Do any types of human disturbances stand out in terms of their effects on contemporary rates of phenotypic change (Pelletier & Coltman, 2018). We envision this last analysis as a precursor to what will surely be additional analyses of the new (and future) dataset with current and future statistical approaches. To synthesize, we propose a new perspective on how to study contemporary rates of change in future studies.

## METHODS & RESULTS

### Database development

The current dataset has a series of notable changes relative to earlier versions of the dataset. First, we added new data that met the necessary criteria (phenotypic traits were quantified from natural populations of the same species either at two time points in the same populations or in two populations with known divergence time) (Fig. S1), and recorded meta-data as described in Box 1. Considering the important number of studies available, it was not possible for us to incorporate *all* studies where rates of changes could possibly be incorporated. We thus, added articles that came to our attentions while also conducting systematic searches in Google Scholar and Web of Science. An important addition to the database was data from 1072 salmonid populations from four species (Clark, Brenner, & Lewis, 2018; Oke et al., 2020). Second, we modified and expanded the categorization of types of human disturbances as defined in Box 2. Third, we checked both old and new data entries to fix any errors. These efforts were facilitated by – whenever possible – obtaining summary data (means, sample sizes, and standard deviations) directly from tables, figures, or by contacting authors, from which we calculated rates of change (details below). Total number of rates for each disturbance type by study design (allochronic/synchronic), type of study (genetic/phenotypic), and taxa are presented in Table S1.

### The data

Here we outline processes common to all analyses. First, all statistical analyses were performed in R environment 4.0.5 (R Core Team, 2021). Second, we only include studies in which the number of generations elapsed was 300 or fewer (7338 rates), which is suitable for analyses of “contemporary” change (Hendry et al., 2008). Third, most analyses were conducted using both darwins and haldanes (each in separate analyses, never combined) – because the two metrics have different biological and statistical properties, as well as different data requirements (Gingerich, 1993; Hendry & Kinnison, 1999; Hunt, 2012). Darwins are defined as the proportional change in the mean trait value in units of *e* per million years and are appropriate for data on a ratio scale, but not an interval scale (Box 1). We have 287 rates on interval scale from which we could not calculate Darwins. Darwins were calculated as

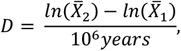

where 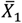 and 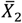 are either mean trait values for one population at times 1 and 2 or mean trait values for two populations that had a common ancestor at a known time in the past, which then scales “time” as million years in the denominator. Haldanes are the change in the mean trait value in standard deviations per generation. Haldanes were calculated as

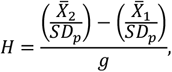

where 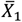 and 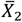 are the mean trait values as explained above, SD_p_ is the pooled standard deviation of the two samples, and *g* is the elapsed time in generations (i.e., number of years divided by generation length (Box 1)).

The response variable in all subsequent analyses was the numerator of the rate metric (i.e., darwin or haldane numerator). To avoid self-correlation when plotting darwins and haldanes against time intervals, we use the absolute amount of change (numerators) against time intervals (Hendry & Kinnison, 1999). For all analyses (except questions 3 and 4) we used the mean amount of phenotypic change for a given species/system/study (Box 1) to avoid nonindependence of data points within a system (Hendry & Kinnison, 2001). Finally, we calculated effect sizes using partial η^2^ to compare the original studies to the updated ones. We note that partial η^2^ represents the proportion of variation in a particular response variable that is explained by predictor variables and is therefore sensitive to the total amount a variance in a given dataset. We now present analyses specific to each question, their respective results, and a brief discussion of results.

## THE QUESTIONS

### Question I: Are rates of phenotypic change greater in human-disturbed systems?

Humans cause particularly dramatic environmental changes, and so we might expect human disturbance to accelerate rates of phenotypic change. Consistent with this idea, Hendry et al., (2008) reported that phenotypic rates of change were higher for populations in human-disturbed systems than for populations in more “natural” systems that were not subject to direct human disturbance. To replicate the original analyses with our new dataset, we used analyses of covariance (ANCOVA) to assess whether the absolute amount of phenotypic change (darwin or haldane numerator) differed between the two general contexts (human-disturbed or natural) while controlling for the length of the time interval (years for darwins, generations for haldanes). We ran two separate analyses: one for haldane numerators and one for darwin numerators.

As in Hendry et al., (2008), our new dataset suggests that rates of change were generally higher in human-disturbed systems compared to natural systems (Fig. 1; Fig. S2). The difference between contexts (human-disturbed versus natural) was, however, reduced in our new dataset, partial η^2^ = 0.018, compared to the original analysis, partial η^2^ = 0.115, (Fig. 1; Fig. S2; Table 1; Table S2). This smaller difference could result from confounding effects of multiple types of disturbances (Galton, 1886; Kelly & Price, 2017; Pelletier & Coltman, 2018). That is, it can be difficult to assign a single type of human disturbance to a particular system. For example, climate change is likely to affect all systems indirectly, including systems we have classified as “natural.” Here, we classified disturbance as climate change only if the original study specifically tested for an effect of climate change. We will later return to the influence of these multiple disturbances on inferences about contemporary evolution. Our expanded dataset could also mean that effects like winnowing, which would reduce the number of populations with low rates of phenotypic change because they are more likely to go extinct (Hendry et al., 2008), do not appear to have a strong overall effect.

**Table 1.** Partial η^2^ values for darwin numerators and their appropriate models for each study revisited. Type refers to if the data were from a phenotypic study or a genotypic study (e.g., common garden). For *p*-values for question V., see Table 4.

| Question | Model | N<br>original | N<br>updated | Explanatory<br>variable | Effect size<br>original | Effect size<br>updated | $p$<br>original | $p$<br>updated |
| --- | --- | --- | --- | --- | --- | --- | --- | --- |
| I | Absolute Darwin<br>numerators ~ years *<br>human influence | 2844 | 6957 | Years | 0.004 | 0.006 | 0.543 | 0.562 |
|  |  |  |  | Human<br>Influence | 0.115 | 0.018 | **0.031** | **< 0.001** |
| II | Absolute Darwin<br>numerators (means) ~<br>human influence +<br>years | 87 | 1529 | Years | 0.010 | 0.002 | 0.366 | 0.096 |
|  |  |  |  | Human<br>Influence | 0.180 | 0.010 | **< 0.001** | **< 0.001** |
| III | Absolute Darwin<br>numerators ~<br>introductions * years | 104 | 276 | Years | 0.612 | 0.162 | 0.954 | 0.750 |
|  |  |  |  | Introductions | 0.822 | 0.920 | 0.480 | 0.430 |
| V | Log absolute Darwin<br>numerators ~<br>disturbance + type +<br>years +generations |  | 2109 | Years |  | 0.959 |  |  |
|  |  |  |  | Disturbance |  | 0.998 |  |  |
|  |  |  |  | Type |  | 0.041 |  |  |

**Figure 1.**
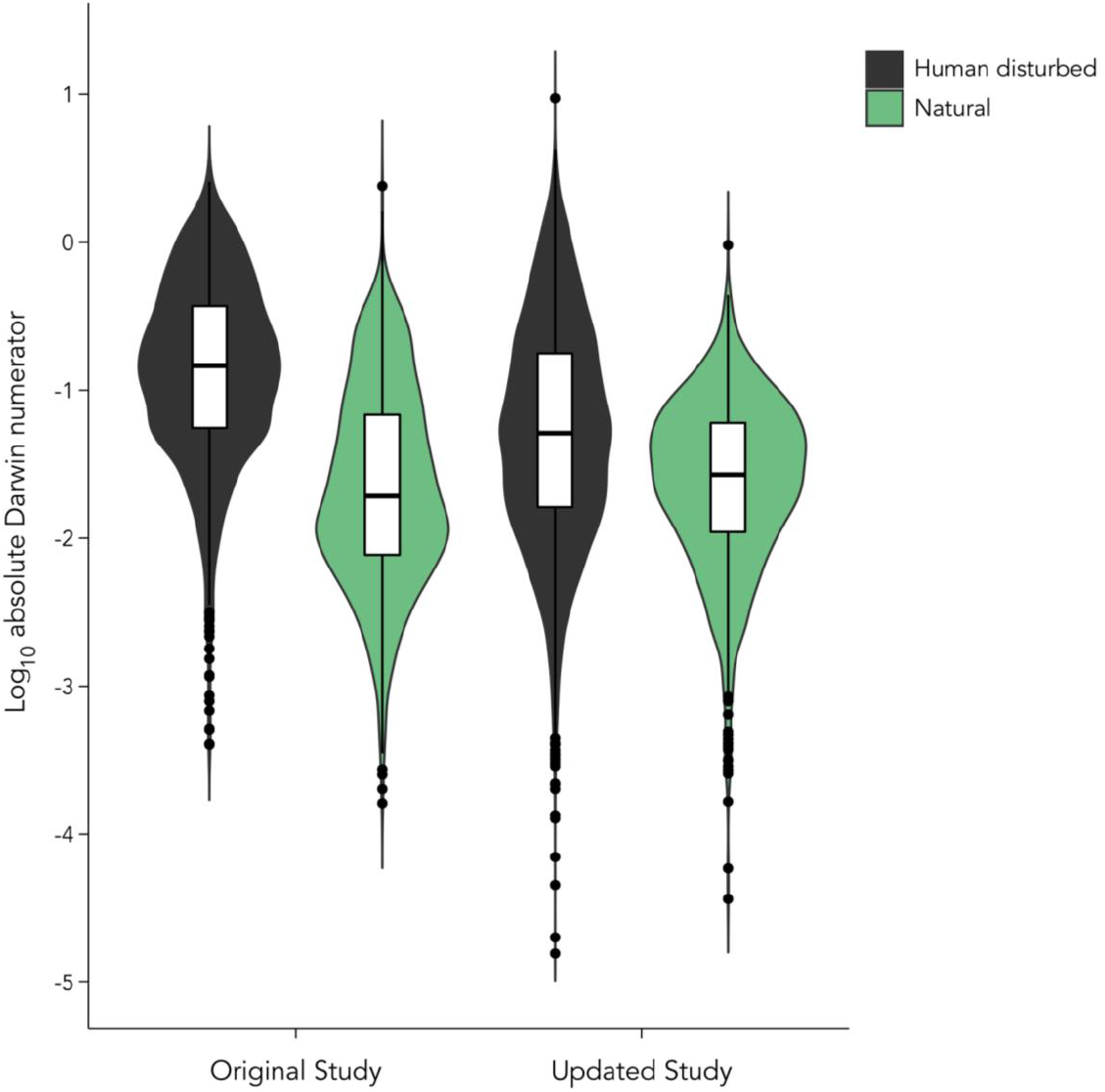
Violin plot of rates of phenotypic change (log-transformed darwin numerator) comparing natural (green) and human disturbed (black) systems. The two left violins are data from Hendry et al., (2008) and the two right violins are from our updated dataset. Axes were cut for better visualization (2 outliers removed from figure).

By comparing estimates from wild populations versus those from common-garden or animal model analyses (Box 1), Hendry et al., (2008) concluded that plasticity likely contributed substantially to the rate differences between human-disturbed and more natural contexts (Fig. S3; Fig. S4). This suggestion arose because the difference between contexts was lower when common-garden or animal model studies were used – and because large changes could sometimes be seen immediately after a disturbance (Hendry et al., 2008). Such patterns also occurred in the present study supporting those original inferences (Fig. S3; Fig. S4). At the same time, it is important to note that genetic changes definitely occurred in a number of studies, but that many of the most disturbed contexts (e.g., harvesting) are not particularly amenable to the assessment of genetically-based phenotypic change. Only 24% of the phenotypic change entries in our dataset could be labelled as genetically based, provided an adequate experimental design.

### Question II: Are particularly rapid and consistent changes associated with harvesting?

A particularly strong and consistent disturbance that directly impacts some populations occurs when humans act as a predator, such as in cases of harvesting. To explore this idea, Darimont et al., (2009) took the human-disturbed versus natural distinction of Hendry et al., (2008) and divided the human-disturbed systems into those experiencing direct harvesting versus those experiencing other forms of human disturbances. After adding more data, especially to the harvesting category, Darimont et al., (2009) reported that populations subject to harvesting had increased phenotypic rates of change compared to other types of human disturbance and also compared to natural systems. Here, we only use darwins (not haldanes) as to replicate the original analysis. To replicate that original analysis with our updated dataset, we used ANCOVA to compare darwin numerators associated with harvesting to those associated with other types of human disturbances (combined) and those associated with more natural systems – while including time (years) as a covariate. We ran ANCOVAs using both mean and maximum rates of change per system to replicate the original study.

As in Darimont et al., (2009), our new data suggested that phenotypic changes associated with harvesting are greater than those associated with other types of human disturbances or natural systems (Fig. 2; Fig S5; Table 1). However, the effect size (partial η^2^) of human disturbance (three contexts: harvesting, other human disturbance, or natural) decreased from 0.180 in the original study to 0.010 in the updated dataset (Table 1). We suggest that this decrease in effect size is mainly driven by large datasets containing many study systems with high variation. As an example, when we remove the Oke et al., (2020) and Clark et al., (2018) datasets, the effect size of human disturbance increases to 0.139 (Fig. S6). These datasets correspond to salmonid data for 1072 populations from four species over a time frame of up to 77 years. Of these populations, 60% are (or were) likely subject to harvest – and so needed to be added to that category in the present analysis. The remaining 40% of populations were added to the natural category. However, the strength and type of harvest is expected to be highly variable among those species and populations. We note that this variability in harvest strength is likely true for most harvested populations. This issue again highlights the difficulty of unambiguously assigning human disturbances to systems that are surely experiencing multiple types of disturbance that vary in type and intensity across systems.

**Figure 2.**
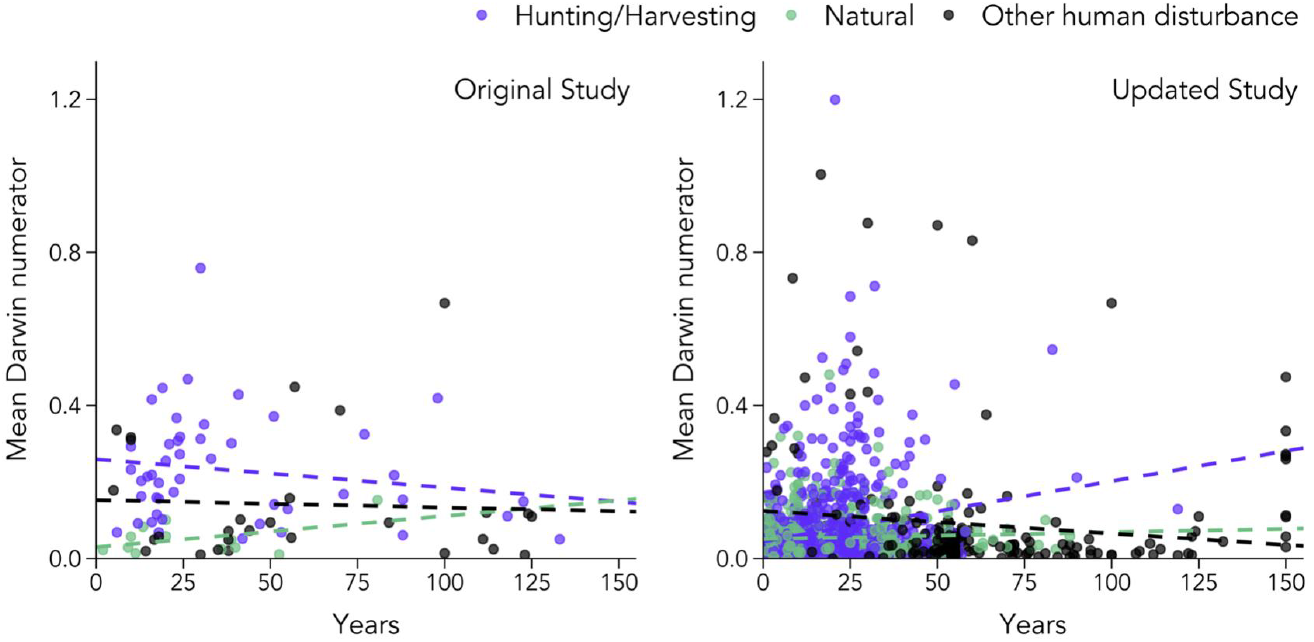
Rates of phenotypic change (mean darwin numerator) for hunted/harvested systems (purple), natural systems (green), and other types of human disturbed systems (black). Rates of change are averaged values per study system. Left panel is data from Darimont et al., (2009) and the right panel is our current dataset.

Despite the lower effect size for harvesting in our new dataset, we note that harvested systems represented 35 of the 50 largest mean phenotypic changes in the dataset. Hence, it seems likely that humans as predators generate a diversity of rates of change – from many slow rates to some exceptionally high rates. Our findings continue to support the claims of fisheries and hunting wildlife scientists who have long argued the lasting effects of harvesting on some (but not all) natural populations (Kuparinen & Festa-Bianchet, 2017; Morrissey, Hubbs, & Festa-Bianchet, 2021; Pigeon, Festa-Bianchet, Coltman, & Pelletier, 2016; Van de Walle, Pigeon, Zedrosser, Swenson, & Pelletier, 2018).

### Question III: Do introduced populations show particularly rapid rates of change?

When a species is introduced into a new location, it often experiences massive shifts in biotic and abiotic conditions, which are expected to cause particularly rapid phenotypic changes (Carroll, 2007; Cox, 2004; Huey, Gilchrist, & Hendry, 2005; Kinnison, Unwin, & Quinn, 2008). However, Westley (2011) used an earlier version of this dataset to report that introduced populations do not – in fact – evince particularly strong or consistent phenotypic changes when compared to non-introduced populations. To replicate their analyses, we first averaged rates by species and then used ANCOVA to compare the magnitude of phenotypic change (darwin or haldane numerators) between introduced (disturbance classified as introduced) and non-introduced (all other disturbance categories) populations with time (years or generations) as a covariate (Westley, 2011).

We found that, on average, introduced populations have higher rates of change than non-introduced populations (Fig. 3; Fig. S7). Although the difference between introduced and non-introduced populations is marginal, the addition of our new data increased the effect size (Table 1). This finding is consistent with the evidence that introduced populations do sometimes show very rapid rates of change (Fig. 3). For example, zebra mussels (*Dreissena polymorpha*) introduced in European lakes rapidly shifted their growth rates (Czarnołeski et al., 2005) and Eastern grey kangaroos, (*Macropus giganteus*) introduced from Tasmania to Maria Island rapidly shifted their behaviour when no longer subject to predation (Blumstein & Daniel, 2003). This is consistent with Westley’s conclusion that a small number of introduced species having very high rates of change drove the perception that introduced species show rapid change. Indeed, the updated dataset has both very high rates and very low rates of change in introduced populations (Fig. 3).

**Figure 3.**
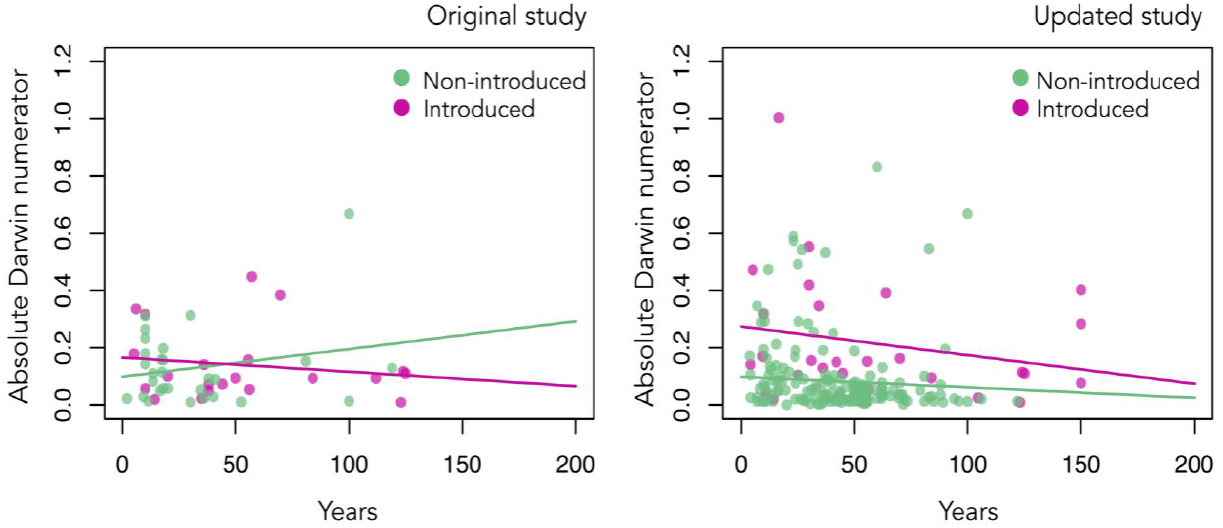
Absolute phenotypic change in introduced species (pink) and non-introduced species (green) measured in absolute darwin numerators. Each point is an expression at the species taxonomic level. Darwin numerators are plotted as a function of years. The left panel is data from Westley (2011) and the right panel is our updated dataset.

Populations introduced to a new habitat likely experience abrupt directional selection that drives rapid evolutionary rates compared to non-introduced populations (Carroll, 2007; Cox, 2004). Once a population achieves adaptive optima and is more locally adapted, phenotypic rates of change are expected to decline with time (Fig. 3). This expectation is consistent with the declining rates of change with time since introduction (Figs. 1 & 2 in Westley 2011), although that pattern is also consistent with other processes (Hendry & Kinnison, 2001). These processes include averaged rates over longer time spans or depletion of genetic variation (Kinnison & Hairston, 2007). Finally, we note that both introduced and non-introduced populations are also experiencing other types of human disturbances that can influence their rates of evolution, thus potentially obscuring effects of introduction per se.

### Question IV: Are body sizes increasing through time as suggested by Cope’s Rule?

Cope’s Rule states that lineages generally evolve larger body sizes over evolutionary time (Cope, 1885). Based on an analysis of selection estimates, Kingsolver and Pfennig (2004) argued that evidence of directional selection for larger body size in contemporary populations is consistent with a microevolutionary explanation for Cope’s Rule. However, only a few studies support the trend suggested by Cope’s Rule (Baker, Meade, Pagel, & Venditti, 2015; Siepielski et al., 2019). In fact, when looking for evidence of general trends toward larger mean body size in contemporary populations, Gotanda et al., (2015) found no such trend and instead found a general trend towards decreasing body size.

To replicate the analysis of Gotanda et al., (2015), we used the raw (i.e., signed, rather than absolute value) estimates of haldane or darwin numerators for allochronic studies (Box 1). The original analyses did not average per system or species, and so all new analyses are similarly based on individual rates. We first square-root transformed 2-D traits (e.g., surface area) and cube-root transformed 3-D traits (e.g., volume or mass) and then compared rates for body size traits to rates for other types of traits in a one-tailed Wilcoxon rank-sum test (see Gotanda et al., 2015 for details). We next conducted a sign test to determine whether the change in body size within populations was more commonly positive or negative. Finally, we re-ran the sign tests excluding rates calculated from populations known to be subject to harvesting – as harvesting is expected to cause particularly rapid decreases in body size (Darimont et al., 2009; Sharpe & Hendry, 2009; see question 2).

When compared to other traits, rates of change for body size are not larger (Table 2; Fig S8). As in the original analysis of Gotanda et al., (2015), rates of change in body size are more often negative (organisms are getting smaller overall), even when excluding populations subject to harvesting (Table 3; Fig. 4; Fig S9). Further, rates of change for body size are not more positive (or less negative) when compared to other traits (Table 3; Fig. S9), and the type of human disturbance does not appear to affect rates of change on body size (Fig. 4). These results are matched by other recent analyses of body size trends in a variety of taxa (Gardner, Peters, Kearney, Joseph, & Heinsohn, 2011; Sheridan & Bickford, 2011). For instance, recent research suggests that body sizes are broadly decreasing as a response to climate change perhaps due to effects of increasing temperature and variable precipitation on organismal development and growth (Fryxell et al., 2020; Sheridan & Bickford, 2011). Based on our findings, and previous findings, we emphasize that Cope’s Rule is not a general rule, and rather a trend seen in certain groups of organisms (Baker et al., 2015; Rollinson & Rowe, 2015; Waller & Svensson, 2017).

**Table 2.**
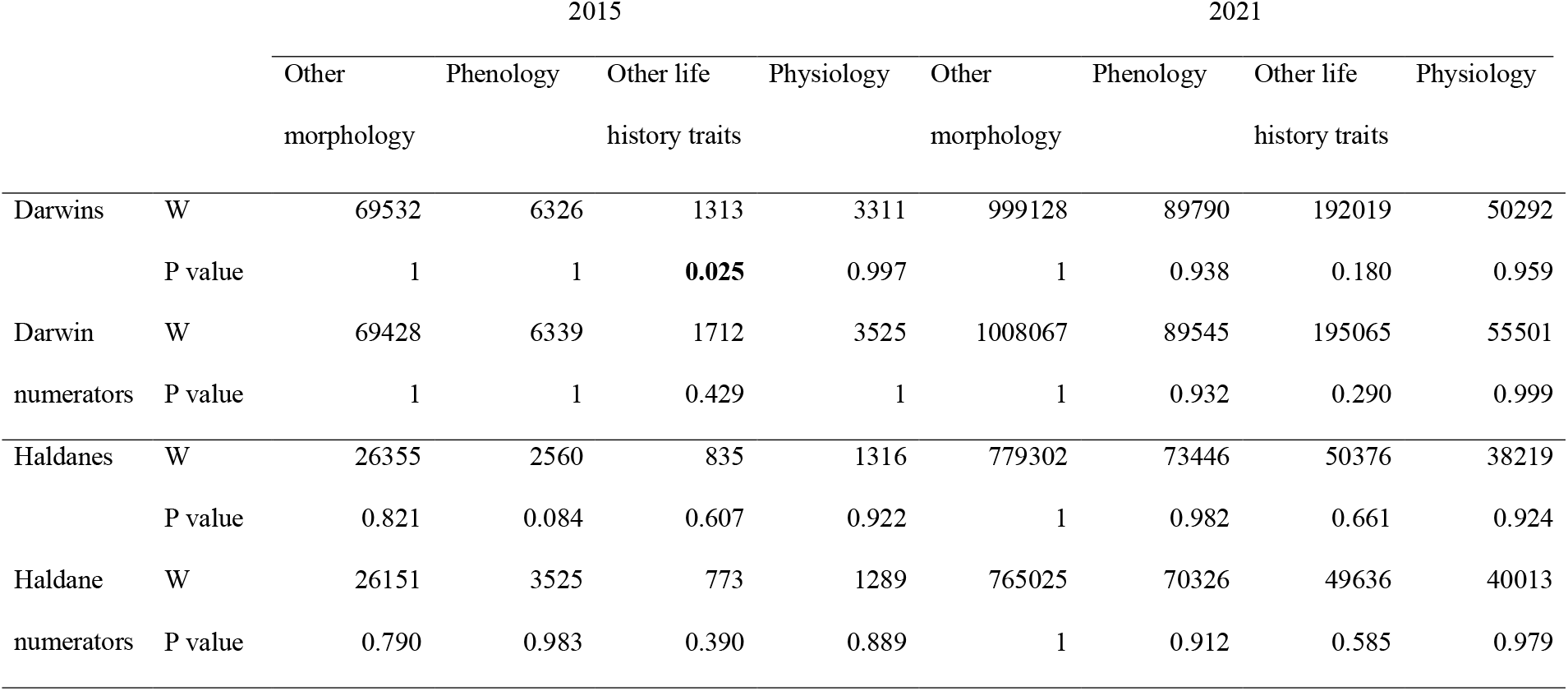
Pairwise Wilcoxon signed-rank test results for size *versus* a different phenotypic trait (one-sided) to see if rates of evolution for body size were higher than other traits. Bold indicates significant P values where body size rates are higher than the other phenotypic trait, though not necessarily *positive*. Trait classification followed the definitions found in Kingsolver and Diamond (2011).

**Table 3.**
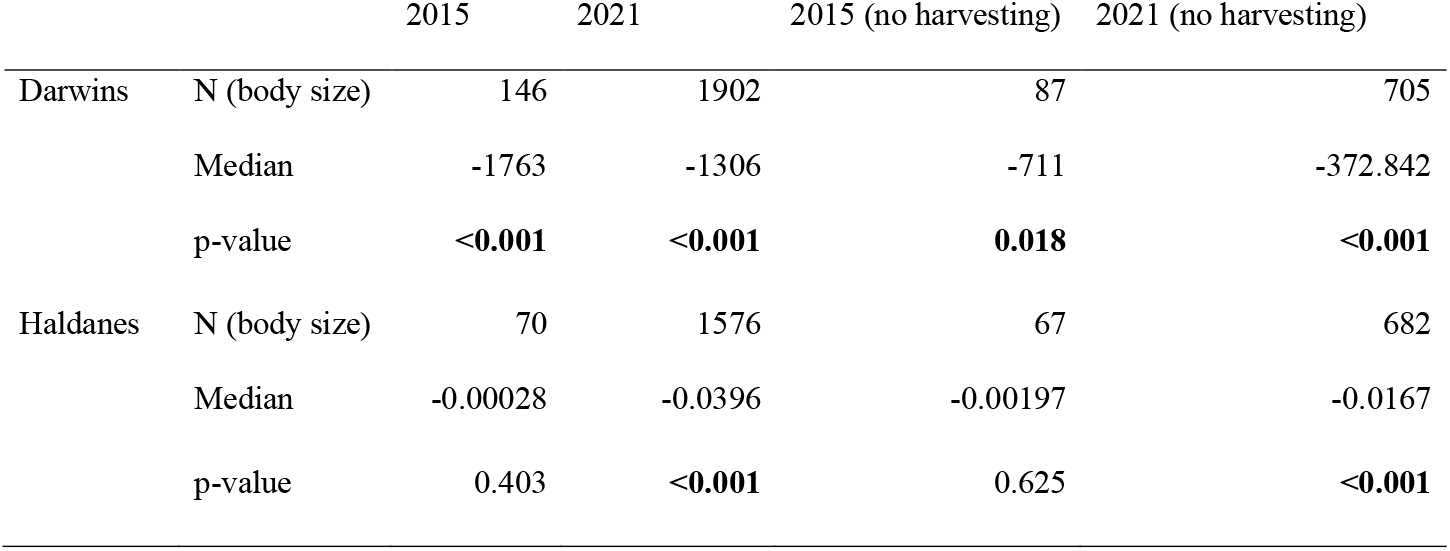
Sign-test results for rates of evolution testing whether body size rates were significantly different from zero. Results are shown for all data and for data with harvesting data removed. Median rates are given, and bold values mean the median is significantly different from zero. Due to the nature of the sign test, numerators yield the exact same results, and so are not reported. Body size classification followed the trait classification definitions found in Kingsolver and Diamond (2011).

|  |  | 2015 | 2021 | 2015 (no harvesting) | 2021 (no harvesting) |
| --- | --- | --- | --- | --- | --- |
| Darwins | N (body size) | 146 | 1902 | 87 | 705 |
|  | Median | -1763 | -1306 | -711 | -372.842 |
|  | p-value | **<0.001** | **<0.001** | **0.018** | **<0.001** |
| Haldanes | N (body size) | 70 | 1576 | 67 | 682 |
|  | Median | -0.00028 | -0.0396 | -0.00197 | -0.0167 |
|  | p-value | 0.403 | **<0.001** | 0.625 | **<0.001** |

**Figure 4.**
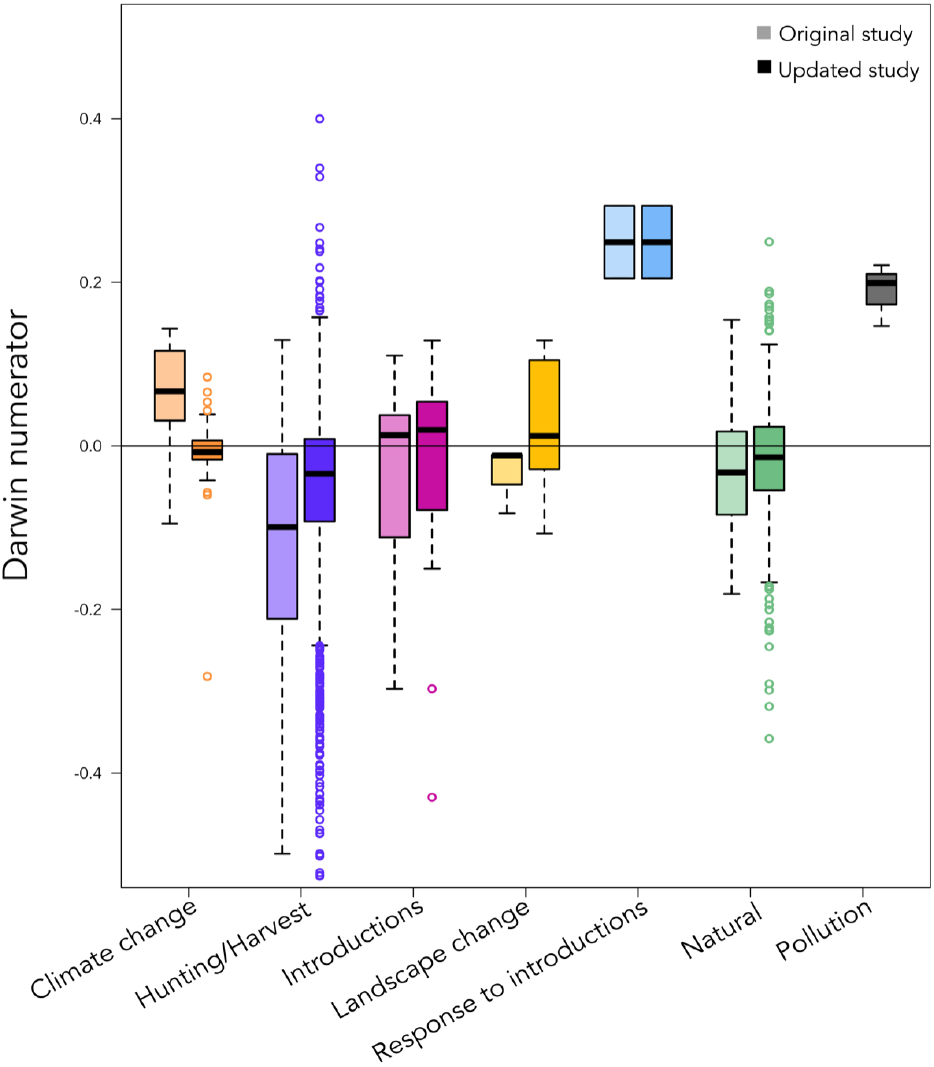
Boxplots depicting rates of change associated with body size measured in darwin numerators for each disturbance category (climate change, hunting/harvesting, introduction, landscape change, response to introductions, natural, and pollution). Light boxes are results from Gotanda et al., (2015) and dark boxes are results from our updated dataset. Y axis was truncated to aid in visual assessment.

### Question V: Does any type of disturbance stand out with respect to rates of change?

Based on the previous studies we have now revisited, as well as more recent reviews (Pelletier & Coltman, 2018), we expect associations between high rates of phenotypic change and particular types of human disturbances such as pollution (Hamilton, Rolshausen, Uren Webster, & Tyler, 2017), harvesting (Sullivan, Bird, & Perry, 2017), or landscape change (Legrand et al., 2017). Using our extensive dataset, we are now able to ask if these or other types of human disturbances stand out with respect to rates of phenotypic change.

To answer this question, we used more advanced analyses – in contrast to the above questions. We used general linear models in which the response variables were log10-transformed absolute darwin (or haldane) numerators, and independent variables included type of human disturbance, time (years or generations, log10-transformed), and type of study (genetic or phenotypic). Darwin and haldane numerators were analysed in separate analyses. As in question 4, we first square-root transformed 2-D traits (e.g., surface area) and cube-root transformed 3-D traits (e.g., volume or mass) before log transforming the rates. Finally, we used Tukey post-hoc tests (Hothorn, Bretz, & Westfall, 2008) to explore the differential effects of disturbances on differences in rates of evolution.

This new analysis suggests that time (years) had a significant positive impact on darwin rates of change, but type of study (genetic vs. phenotypic) did not (Table 4). The analysis using haldanes suggests similar results where time (generations) had a significant positive impact on haldanes, but not study type (Table S3). We also found that systems associated with pollution have the highest rates of change and that systems associated with climate change have the slowest rates of change (Fig. 5; Fig S10). This finding might be explained by underlying genetic architecture where we expect evolution to occur faster when selection is on one (or a few) loci versus polygenic traits (Kardos & Luikart, 2021; Oomen, Kuparinen, & Hutchings, 2020). In fact, many famous examples of rapid evolution are known to be through selection on single loci such as heavy metal tolerance in plants (Macnair, 1991). Within our dataset, most systems looking at the impact of pollution were plants, which might or might not influence the high rates for this type of disturbance. In short, more work needs to be done to explore potential interactive effects of disturbance type and taxonomic group on rates of change.

**Table 4.**
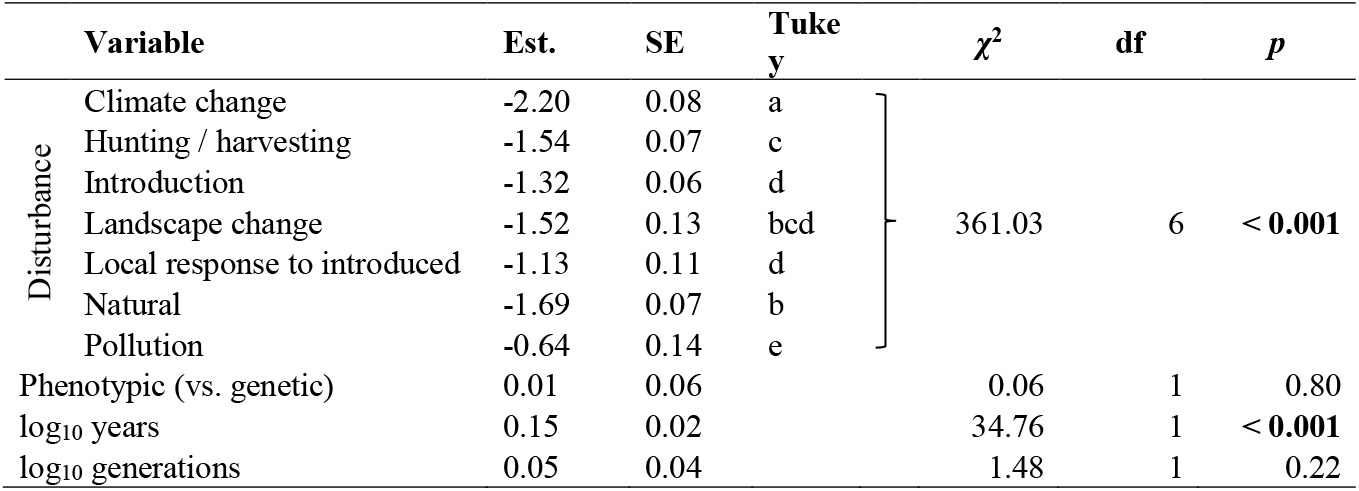
Estimates, standard errors, Tukey test categorizations, and type II likelihood ratio test results for models predicting log_10_ absolute value darwins. Chi-squared values are for the variable-specific likelihood ratio tests.

**Figure 5.**
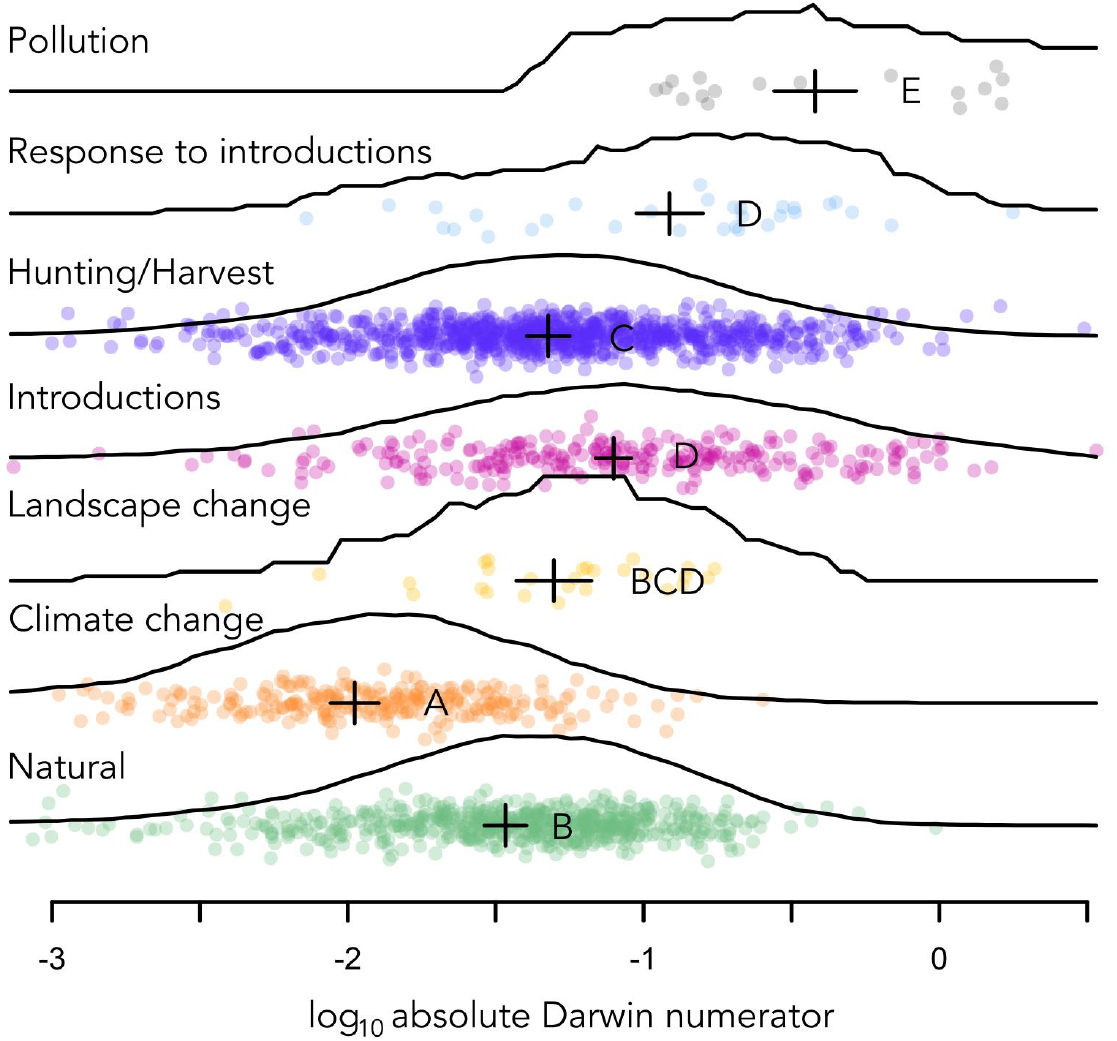
Rates of evolution—in log-transformed absolute darwin numerators—for six types of disturbances (pollution, response to introductions, hunting/harvest, introductions, landscape change, climate change) and natural populations. Points show individual data, lines show smoothed data distributions, and crosses show GLM estimates ± standard errors. Letters indicate characterisations based on Tukey HSD tests. Each point represents a system and reference-specific average.

We found that, on average, phenotypic changes associated with climate change were amongst the slowest. We suspect that climate change has broad reaching effects and is especially difficult to assign as a particular sole causal force. Indeed, climate change must be – at some level – influencing all or most systems included in the dataset, regardless of the disturbance category to which we assigned them. Furthermore, studies focusing on trait change in response to climate change are likely to focus on the more gradual aspects of climate change, such as shifting seasonality and temperature increases (Parmesan & Yohe, 2003), rather than abrupt aspects such as heat waves, storms, and droughts. These more gradual aspects of climate change might also have weak multifarious selection rather than strong selection on single genes. More importantly, climate change can be a particularly noisy environmental driver and so, is especially prone to temporal averaging (Hendry & Kinnison, 1999). In fact, few studies in nature follow populations long enough and at a fine enough temporal resolution to detect fast and large phenotypic changes (e.g., Grant & Grant, 2002). For these reasons, we replicate the analyses by including all populations affected by climate change as “natural” (Fig. S11). The results from this analysis confirm that whether we have a disturbance category dedicated to climate change, or if we include climate change with “natural” systems, our conclusions do not change: climate change and natural systems are still amongst the slowest rates of evolution. Regardless, our new analysis provides the most comprehensive hypothesis (Fig. 5) for how various types of disturbance differ in their effects on rates of phenotypic change.

## DISCUSSION

Conclusions from analyses of earlier datasets of contemporary phenotypic change have largely been upheld in analyses of our new dataset that encompasses a much larger breadth of species, traits, and human disturbances. I. Human disturbed systems have slightly larger rates of phenotypic change then do natural systems (Fig. 1). II. Harvesting by humans results in larger rates of change compared to other types of disturbances (Fig. 2). III. Introduced populations have higher rates of change than non-introduced populations (Fig. 3). IV. There is no trend for increasing body size through time (Fig. 4). V. Systems affected by pollution have larger rates of change compared to other types of disturbances (Fig. 5).

Overall, contemporary rates of phenotype change range from very slow to very fast (relative to other rates), with the latter typically gaining the most attention. This pattern is found in other datasets focusing on estimates of selection in natural populations (Kingsolver et al., 2001; Siepielski et al., 2013). Our database includes some striking examples of rapid phenotypic rates of change. For example, horn size in bighorn sheep (*Ovis canadensis*) decreased by 10% over 19 years when targeted by trophy hunters (Pigeon et al., 2016); body length in harvested Chinook salmon (*Oncorhynchus tshawytscha*) in the Yukon River of Alaska has decreased by 5-7% since the late 1970s (Ohlberger et al., 2020); zinc tolerance in tufted hairgrass (*Deschampsia cespitosa*) in zinc-contaminated soils increased by 80% over 26 years (Al-Hiyaly et al., 1990); and total egg count in soapberry bugs (*Jadera haematoloma*) adapting to an introduced host decreased by 8% over 38 years (Carroll, Klassen, & Dingle, 1998). A recent study showed rapid (genetic) evolution of tusklessness in African savanna elephants (*Loxodonta africana)* in response to poaching during the Mozambican civil war (Campbell-Staton et al., 2021). Human influences clearly shape these and many other phenotypic responses; yet high levels of variation around rates of phenotypic change make the most consistent and dramatic changes hard to confirm. In other words, variation in phenotypic change is highly variable, likely due to various human influences but also due to the other processes that this database cannot ascertain.

Our primary goal in the present paper is to make the new database available to any researchers seeking to leverage phenotypic rates of change to answer questions in ecology, evolution, or conservation biology. For instance, we anticipate that researchers will use the new database to answer a series of questions relevant to contemporary evolution and for better understanding and predicting how wild populations will adapt (or not) to human disturbances. We develop some examples of questions left unanswered, or only lightly broached in prior reviews in box 3.

The use of this new database should be accompanied with an understanding of its limits and the resulting caveats of inference.

I. We suspect a strong ascertainment bias where researchers tend to study systems they suspect are strongly affected by human disturbances (e.g., harvested fish populations) or where phenotypic changes are already documented (e.g., phenology). Thus, average effect sizes for a given disturbance might well decrease with the accumulation of more diverse and objective sets of data – as we have shown above with the decrease in average rates of change for harvested systems after we added the 1320 rates from Oke et al., (2020) and Clark et al., (2018) (Fig. S6). These rates associated with body size declines in salmonids were calculated from data collected by the Alaska Department of Fish and Game and collaborators. As such, these rates represent a very broad sampling of populations across Alaska, not a narrow dataset on just the most impacted populations. Another form of ascertainment bias occurs when disturbances cause some populations to go extinct, in which case their rate of phenotypic change cannot be measured (Hendry et al., 2008). It is currently unknown whether this winnowing effect of extinction (Hendry et al., 2008) biases rates upward (i.e., slower-changing populations are more likely to perish) or downward (i.e., faster-changing populations are more likely to perish – likely because they are experiencing more disturbance).
II. Seeking to attribute a particular phenotypic change to a single disturbance (e.g., climate change) is problematic because most populations will be subject to multiple disturbances – and the degree of a given disturbance will vary dramatically among systems. Thus, we encourage future work to consider variation in disturbance intensity (rather than just presence). As examples, one could relate the strength of harvesting on populations (e.g., local catch rates data) to the rate of change in size or age (e.g., Sharpe & Hendry, 2009) or the rate of climate change experienced by populations (e.g., local temperature change) to their specific rate of trait change (e.g., Franks, Sim, & Weis, 2007; Jenni & Kéry, 2003).
III. The genetic and plastic contributions to trait change remain uncertain for most traits in most systems (Merilä & Hendry, 2014). As more studies accumulate, we might be able to profitably analyze only genetically based phenotypic change, such as from common-garden, reciprocal transplants, or animal-model studies. Regardless, we highlight the importance of controlled experiments in combination with phenotypic change in the wild for several reasons: some populations cannot be analyzed with common-garden or animal-model approaches, phenotypes measured in the lab might not be natural, and the “wild” is where organisms interact with their environment (Hendry, 2017).

The study of eco-evolutionary dynamics was born from the recognition of widespread contemporary evolution and the cyclical feedback with ecological processes (Hendry, 2017). With this dataset, we identified some patterns of contemporary evolution (e.g., pollution is the strongest human influence on phenotypic rates of change) and we can use this information to broaden our understanding of eco-evolutionary feedbacks in the wild. We suggest that the next step is to understand how the feedback dynamics implicit to eco-evolutionary dynamics mechanistically shape emergent patterns of contemporary evolution: are some systems more feedback prone and thus have faster or slower rates of evolution? To address such feedback dynamics, we support the development of a comprehensive database which includes not only phenotypic rates of change but also ecological and environmental rates of change and estimates of selection.

Whilst this dataset can be a steppingstone to further our understanding of eco-evolutionary dynamics, we also advocate that this dataset can allow us to move beyond ‘traditional’ ecological and evolutionary patterns and start to consider societal consequences. In the past, the starting point for the study of contemporary evolution was often population dynamics: “*perhaps the greatest contribution that evolutionary rate estimates will ultimately make is an awareness of our own role in the present microevolution of life and cautious consideration of whether populations and species can adapt rapidly enough to forestall the macroevolutionary endpoint of extinction*” (Hendry & Kinnison, 1999). What is now needed is a widespread and formal assessment of how trait changes shape ecological processes such as population dynamics, communities, and ecosystems, as well as the societal consequences such as the health and well-being of people (Des Roches et al., 2018; Hendry, 2017; Hendry, Gotanda, & Svensson, 2017; Palkovacs, Kinnison, Correa, Dalton, & Hendry, 2012; Stange, Barrett, & Hendry, 2021; Fig. 6). For example, we propose that more studies need to empirically study such feedbacks while also assessing the impacts on nature’s contributions to people (NCPs). Oke et al., (2020) provide a compelling example, by comparing the average body size of Alaskan chinook salmon pre-1990 to the average post-2010. The authors estimate that the overall average 8% decrease in body length could – all else being equal – translate into a 16% decrease in number of eggs per female, a 28% decrease in transport of marine-derived phosphorous into freshwater, a 26% reduction in average number of meals provided per fish for people in subsistence communities, and a 21% decrease in price per pound for commercial fishers.

**Figure 6.**
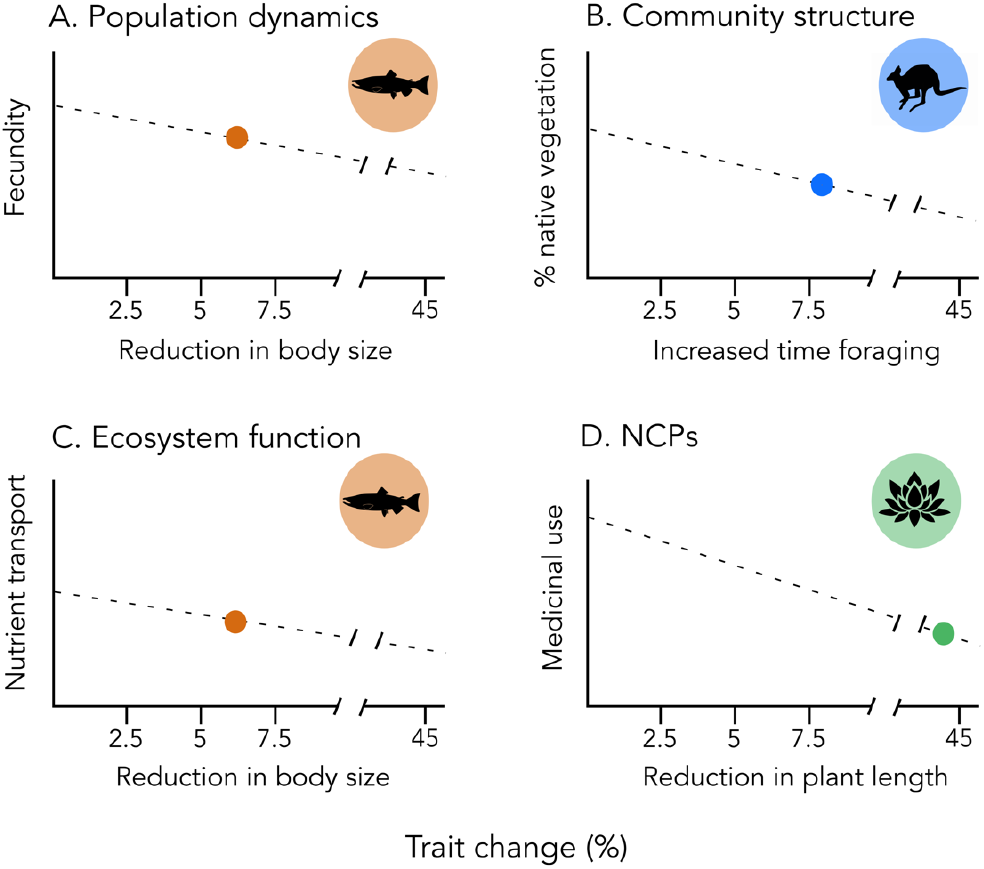
Hypothetical effects of observed phenotypic change for all levels of ecology (populations, communities, ecosystems, and nature’s contribution to people (NCPs)). A) 5-7% decrease in body size in harvested Chinook salmon (*Oncorhynchus tshawytscha*) in the Yukon River (Ohlberger et al., 2020) could decrease fish fecundity, affecting fishery yields. B) 8% increase in time spent foraging in Forester kangaroos (*Macropus giganteus)* introduced to Maria Island (Blumstein & Daniel, 2003) could affect the composition makeup of native vegetation and have effects on ecosystem processes like pollination. C) 5-7% decrease in body size in harvested Chinook salmon in the Yukon River (Ohlberger et al., 2020) could decrease nutrient transport affecting ecosystem properties. D) 43 % decrease in plant length in harvested Himalayan snow lotus (*Saussurea laniceps*) (Law & Salick, 2005) will decrease plant availability for medicinal use by humans.

We would like to close by re-emphasizing that the most dramatic progress will be made through studies that explicitly examine the *consequences* of phenotypic change. That is, more studies should formally calculate the importance of observed (and predicted) phenotypic change for all levels of ecology (populations, communities, ecosystems) and for people (nature’s contributions to people). Determining the genetic and plastic contributions to that change (and its consequences) can then help to determine the limits and opportunities for enhancing or arresting trait changes via conservation and management actions. We here present a hypothetical scenario of such a study using observed rates of change (Fig. 6) where the amount of phenotypic change is correlated with an ecological process that is linked to nature’s contributions to people. The present paper is not the end of an inspiring era of contemporary evolution – it is instead, just the start of future research on not only contemporary evolution, but also on contemporary eco-evolutionary dynamics.

## AKNOWLEDGEMENTS

This work has been supported by NSF OIA 1826777 and NSF OAI 1849227 received by Michael Kinnison. We thank the numerous researchers that shared their data and contributed to the development of The Phenotypic Rates of Change Evolutionary and Ecological Database (PROCEED) Version 5.0. We are especially grateful to both the Carnegie Museum of Natural History and the Powdermill Avian Research Center for sharing their data.

[dataset] Sarah Sanderson, Marc-Olivier Beausoleil, Rose E. O’Dea, Zachary T. Wood, Cristian Correa, Victor Frankel, Lucas D. Gorné,Grant E. Haines, Michael T. Kinnison, Krista B. Oke, Fanie Pelletier, Felipe Pérez-Jvostov, Winer Daniel Reyes-Corral, Yanny Ritchot, Freedom Sorbara, Kiyoko M. Gotanda, and Andrew P. Hendry; 2021; Database reference: Phenotypic Rates of Change Evolutionary and Ecological Database (PROCEED). https://proceeddatabase.weebly.com; Version 5.0

## Box 1.

**Definitions of the different types of data included in the data set**.

1. System: Each system has its own unique species, disturbance, and location (population). Within a given system, you can have multiple traits – for example, tarsus length and fledging date.
2. Disturbance: See Box 2.
3. Trait Classification: Traits were classified as determined by Kingsolver and Diamond 2011: size, other morphology, phenology, other life history traits, behaviour, or physiology.
4. Study type: Studies were determined to be phenotypic in nature if traits were studied in natural populations and studies were determined to be genetic in nature if they used common-garden or quantitative genetic methods (i.e., animal model analyses). We note that studies classified as phenotypic could have a genetic basis (see introduction), but that we could not determine that from the methods of the study.
5. Design: Data were determined to be either allochronic (same population/different time points) or synchronic (populations with known divergence time).
6. Data scale: Data were determined to be ratio (constant interval with a precise zero; e.g., mass or length) or interval (constant interval with an arbitrary zero; e.g., temperature or time of day).
7. Generation time: The amount of time to reproductive age, given in years.

## Box 2.

**Definitions of the different types of human disturbance categories used in the updated database**.

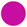 Introductions: “*when humans transferred a species to a new geographical location, and comparisons were then made between introduced and ancestral populations* (Carroll et al., 2005)”.

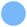 Response to introductions: when a local population of a species is responding to the introduction of a species.

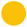 Landscape change: when any type of modification to the habitat of a population occurs.

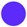 Hunting/Harvesting: when there is hunting or harvesting of a species by humans.

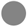 Pollution: when any type of pollutant enters a system.

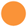 Climate change: when the objectives of the study are directly linked to climate change.

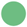 Natural: established populations that are not subject to obvious human impacts (as listed above). Generally, these studies involved the long-term monitoring of natural populations.

## Box 3.

**Questions for future studies using the PROCEED (https://proceeddatabase.weebly.com/)**.

1. Do different organisms or trait types evolve at different rates?
2. Are different disturbances generating confounding or synergistic effects?
3. What are the upper and lower limits to sustainable versus unsustainable evolutionary rates in nature?
4. Does analysing rates of change over shorter timescales (i.e., 5-10) generations, change our inferences? and, do introduced population evolve faster immediately after introductions vs many generations later?
5. Does accounting for error in statistical models change the inferences? (More studies need to publish their errors to make this possible.)
6. Does considering the direction of phenotypic change (positive or negative) change the inferences from previous studies?
7. Analyzing hunting/harvesting records using haldanes when more data become available.
8. Does using different types of effect sizes change the inferences?
9. Is rate of change affected by discrete vs. overlapping generations?

## AUTHOR CONTRIBUTIONS

Ideas: SS, MOB, ROE, VF, MTK, APH, KMG, CC

### Data contributions

SS, MOB, GH, REO, WDR, KBO, KMG, YR, LG, FP, ZTW, APH

### Analyses

SS, MOB, CC, REO, ZTW, KMG

### Writing

SS, KMG, APH, with contributions from all authors

## CONFLIT OF INTEREST

The authors declare no conflict of interest

## DATA AVAILABILITY STATEMENT

The dataset used in this manuscript will be archived on DRYAD and is available at https://proceeddatabase.weebly.com/.

