## Supplemental Materials for "The Pace of Modern Life, Revisited"

### SUPPLEMENTAL INFORMATION

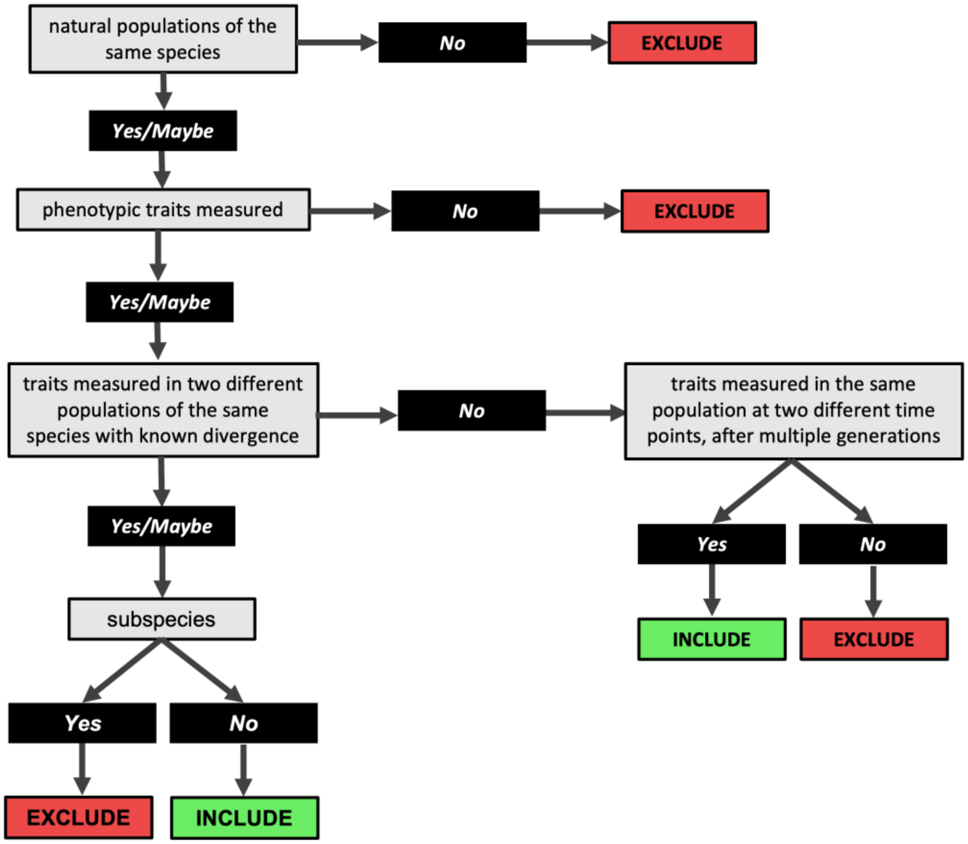


**Figure S1.** Decision tree used when screening abstracts and papers for inclusion in the database.

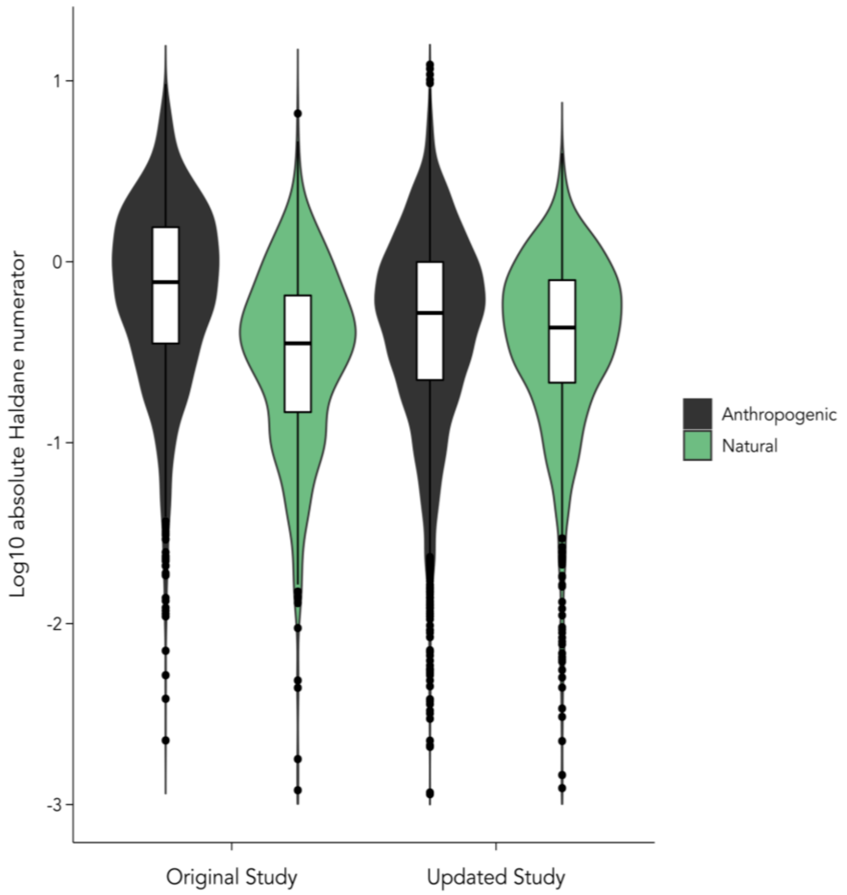


**Figure S2.** Violin plot of rates of phenotypic change (log-transformed haldane numerator) comparing natural (green) and human disturbed (black) systems. The two left violins are data from Hendry et al., (2008) and the two right violins are data from our updated dataset.

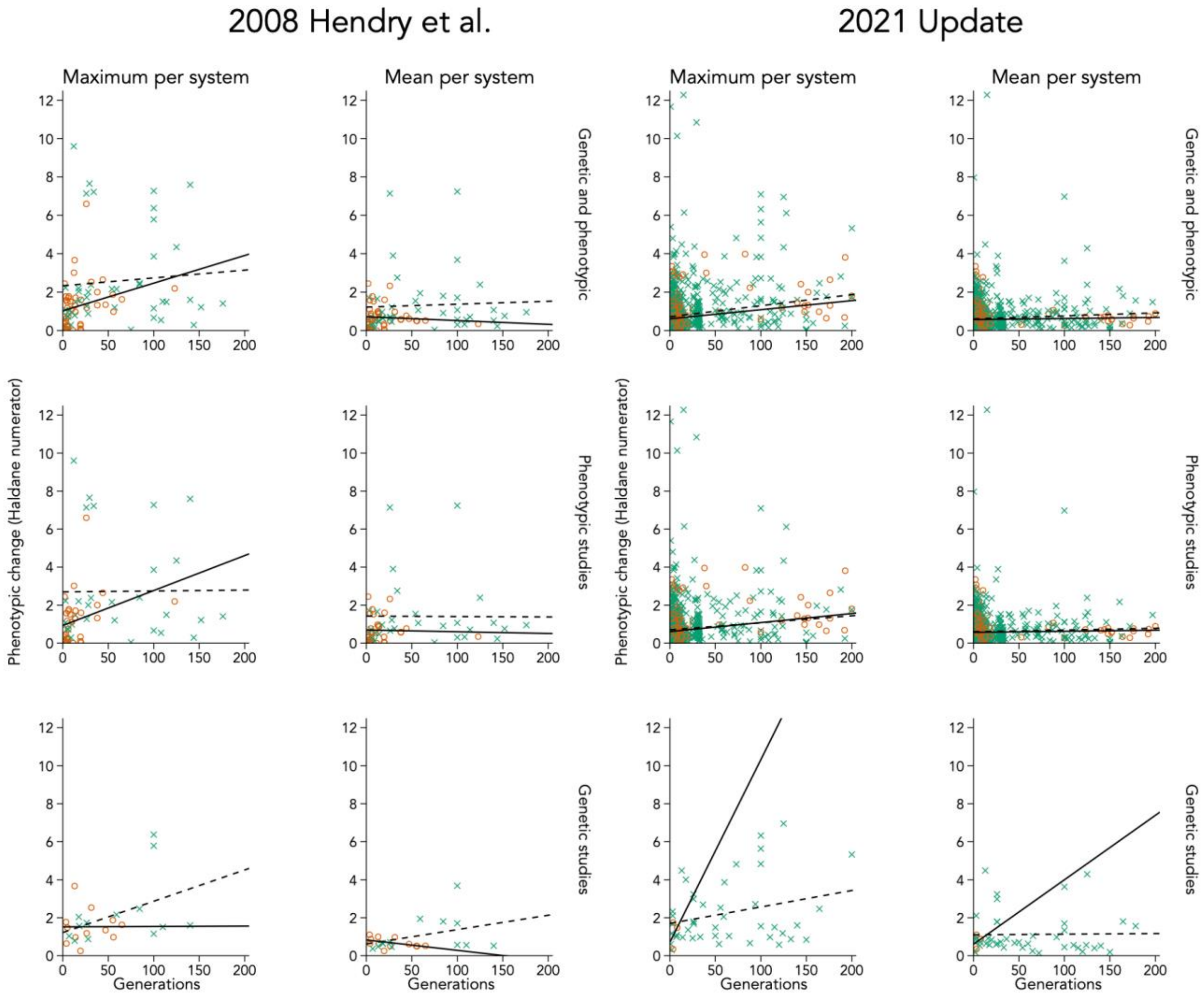


**Figure S3.** Maximum (1st and 3rd columns) and mean (2nd and 4th columns) phenotypic change per system expressed in haldane numerators vs. the time interval (in generations). Top panels show all studies (genetic and phenotypic). Middle panels show studies on wild-caught individuals (phenotypic). Bottom panels show studies using quantitative-genetic studies or common garden experiments (genetic). Changes in human disturbed systems are depicted by crosses and dashes lines whereas natural systems are depicted by open circles and solid lines. Lines are ordinary least-squared regression relationships. The two left columns are data from Hendry et al., (2008) and the two right columns are from the updated dataset.

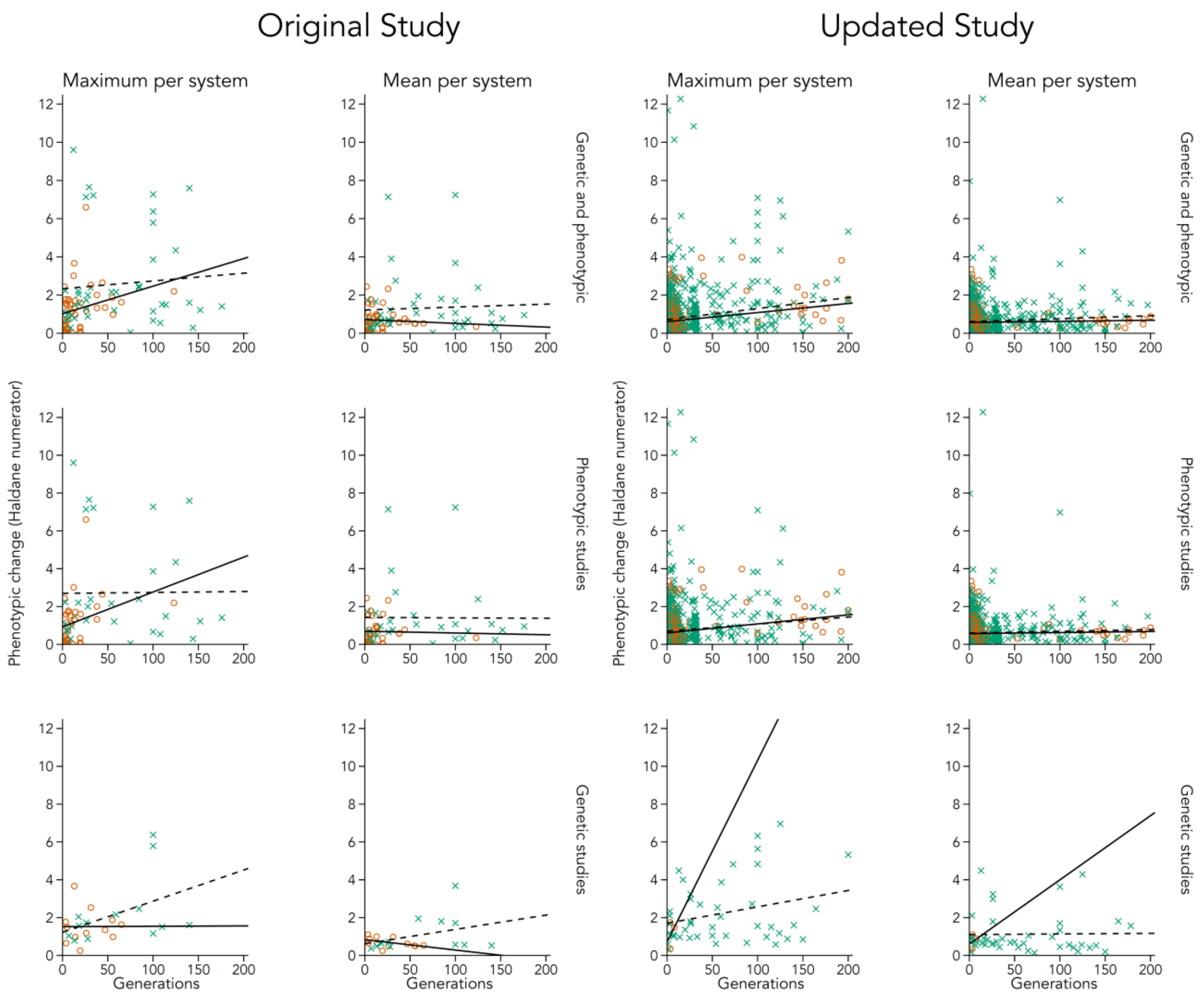


**Figure S4.** Maximum (1st and 3rd columns) and mean (2nd and 4th columns) phenotypic change per system expressed in darwin numerators vs. the time interval (in years). Top panels show all studies (genetic and phenotypic). Middle panels show studies on wild-caught individuals (phenotypic). Bottom panels show studies using quantitative-genetic studies or common garden experiments (genetic). Changes in human disturbed systems are depicted by crosses and dashes lines whereas natural systems are depicted by open circles and solid lines. Lines are least-squared regression relationships. The two left columns are data from Hendry et al., (2008) and the two right columns are from our updated dataset.

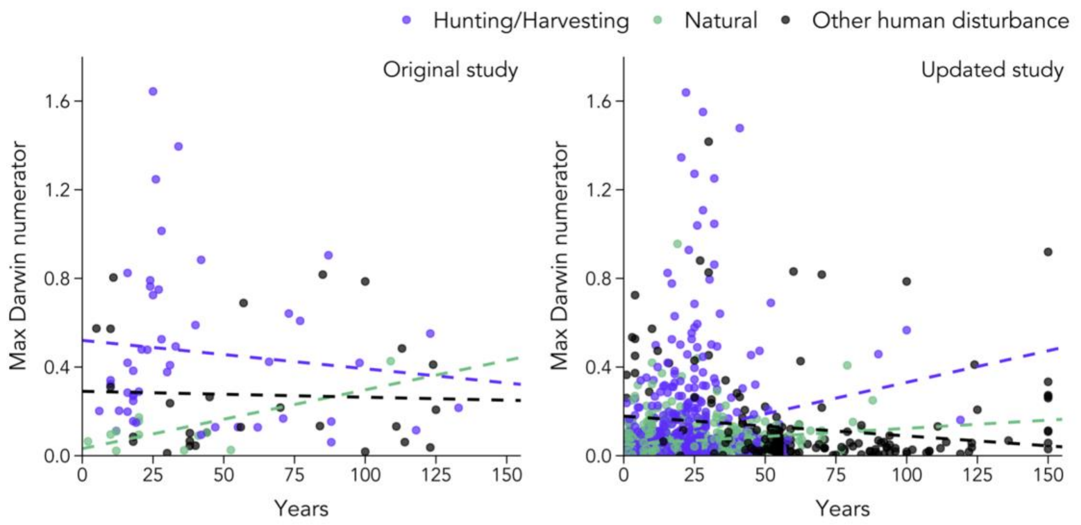


**Figure S5.** Rates of phenotypic change (mean darwin numerator) for hunted/harvested systems (purple), natural systems (green), and other types of human disturbed systems (black). Rates of change are averaged values per study system. Left panel is data from Darimont et al., (2009) and the right panel is our current dataset.

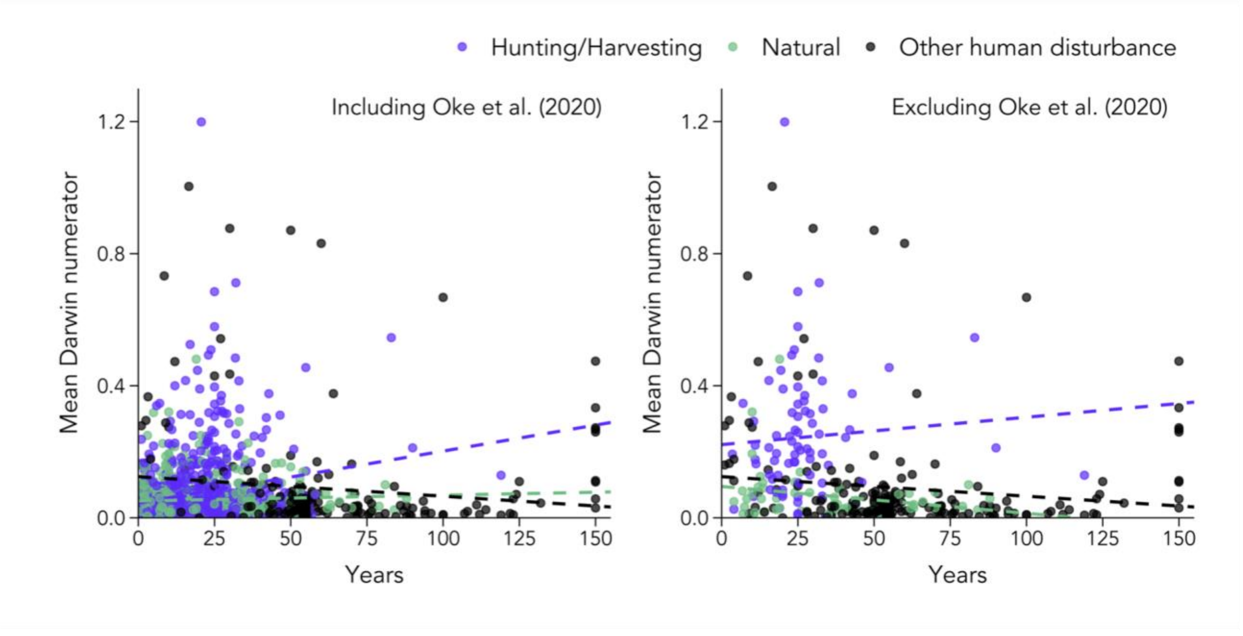


**Figure S6.** Rates of phenotypic change (mean darwin numerator) for hunted/harvested systems (purple), natural systems (green), and other types of human disturbed systems (black). Rates of change are averaged values per study system. Left panel is the entire current dataset and right panel is excluding Oke et al., (2020) data which includes the Clark et al., (2018) dataset.

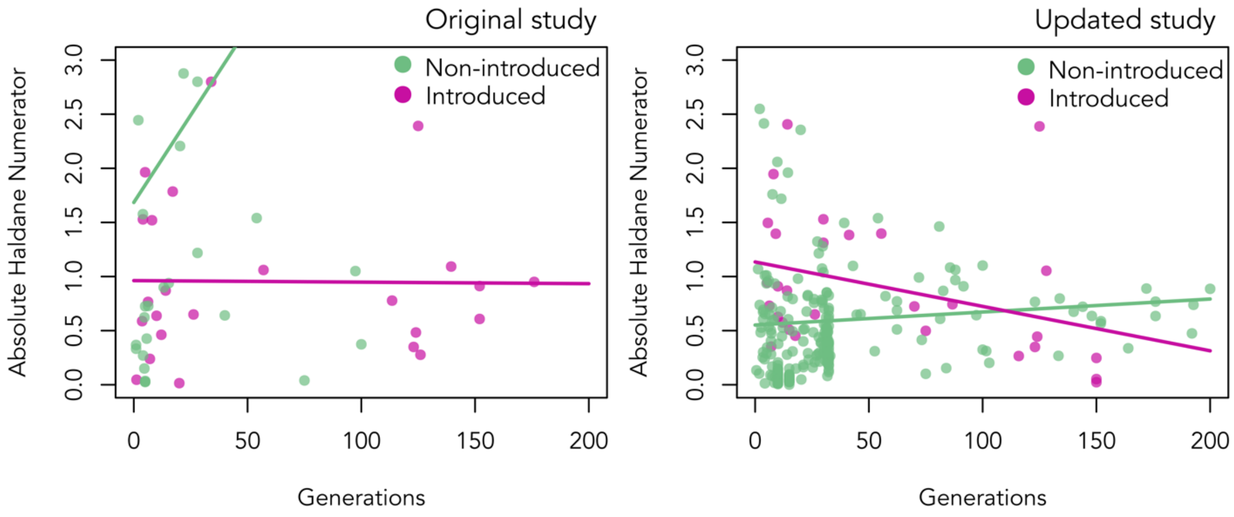


**Figure S7.** Absolute phenotypic change in introduced species (pink) and non-introduced species (green) measured in haldane numerators. Each point is an expression at the species taxonomic level. Haldane numerators are plotted as a function of years. The left panel is data from Westley (2011) and the right panel is our updated dataset.

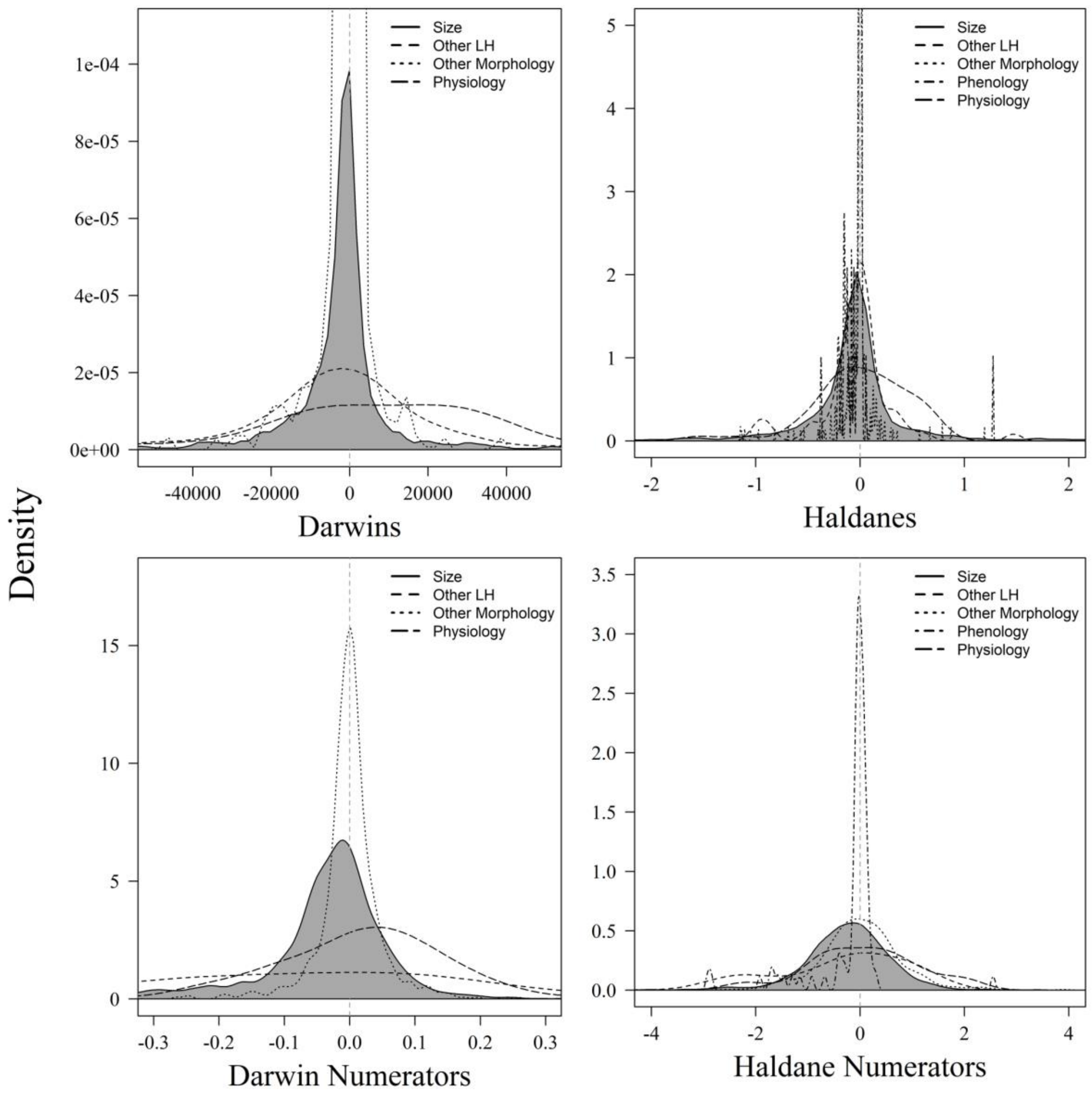


**Figure S8.** Density histograms for darwins, haldanes, and their numerators by five trait classes. Shaded area indicates body size. Y axes are truncated for ease in interpreting the figure.

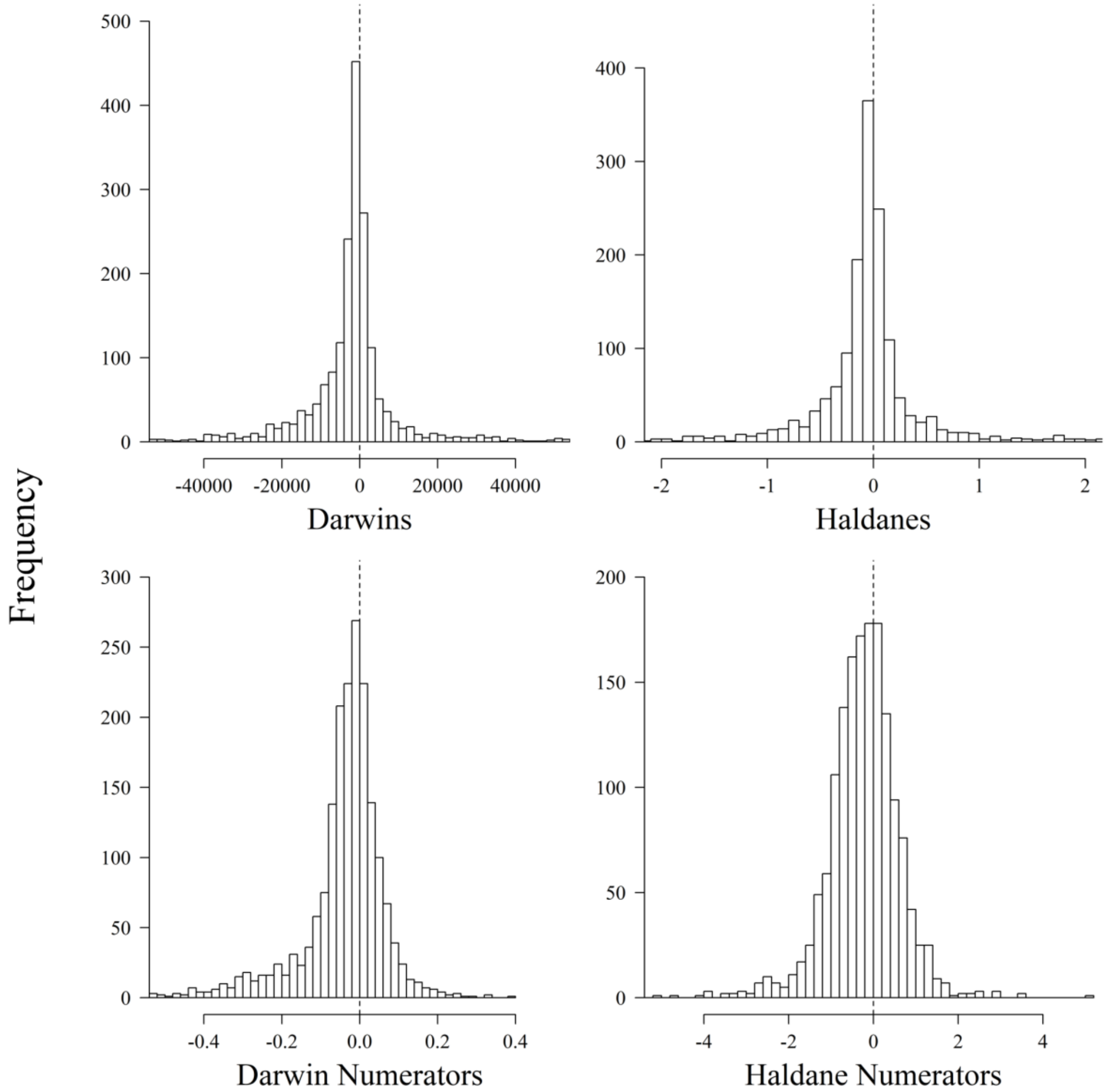


**Figure S9.** Frequency histograms for darwins, haldanes, and their numerators for body size.

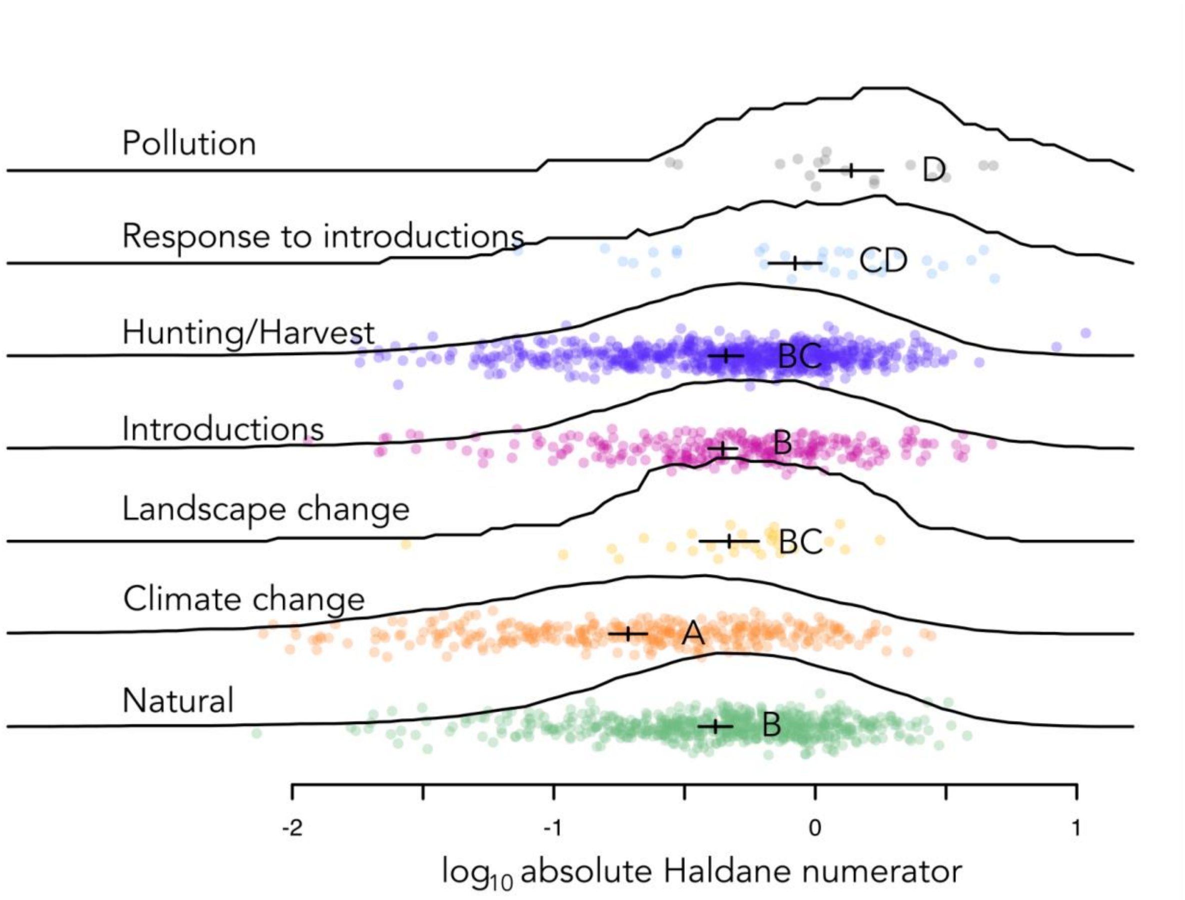


**Figure S10.** Rates of evolution—in log-transformed haldane numerators—for six types of disturbances and natural systems. Points show individual data, lines show smoothed data distributions, and crosses show GLM estimates ± standard errors. Letters indicate characterisations based on Tukey HSD tests. Each point represents a system and reference-specific average.

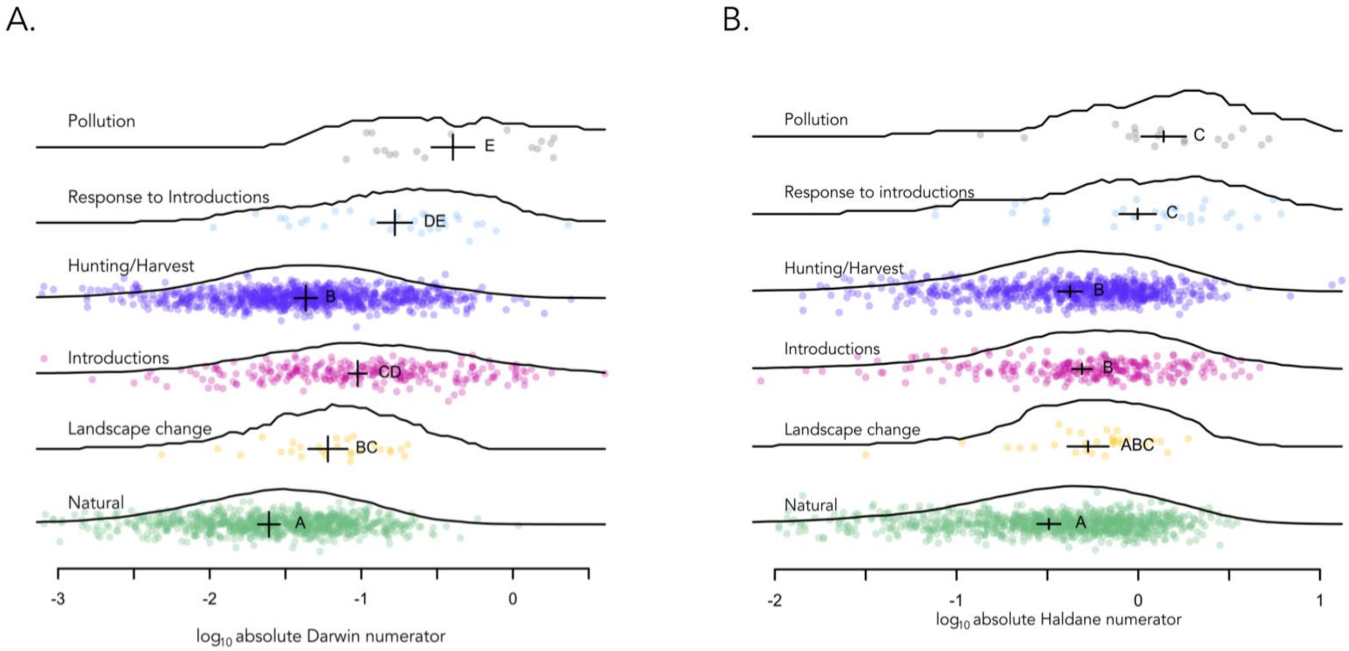


**Figure S11.** Rates of evolution—(A) in log-transformed darwin numerators and (B) log-transformed haldane numerators (B)—for five types of disturbances and natural systems (including climate change). Points show individual data, lines show smoothed data distributions, and crosses show GLM estimates ± standard errors. Letters indicate characterisations based on Tukey HSD tests. Each point represents a system and reference-specific average.

**Table S1.** Total number of rate entries in the database for each type of disturbance by study design, type of study, and taxa. Refer to boxes 1 and 2 for definitions.

| | Study design | | Type of study | | Taxa | | | | | |
|---|---|---|---|---|---|---|---|---|---|---|
| Disturbances | Allochronic | Synchronic | Genetic | Phenotypic | Birds | Fish | Herps | Invertebrates | Mammals | Plants |
| Climate change | 373 | 0 | 11 | 362 | 248 | 0 | 6 | 2 | 111 | 6 |
| Hunting/harvest | 1387 | 1 | 0 | 1388 | 0 | 1322 | 0 | 0 | 33 | 12 |
| Introductions | 90 | 3840 | 1527 | 2403 | 932 | 1557 | 78 | 947 | 104 | 312 |
| Landscape change | 41 | 94 | 0 | 135 | 6 | 128 | 1 | 0 | 0 | 0 |
| Response to introductions | 13 | 193 | 150 | 56 | 0 | 2 | 1 | 148 | 1 | 54 |
| Natural | 1186 | 43 | 13 | 1216 | 263 | 561 | 0 | 33 | 365 | 7 |
| Pollution | 39 | 38 | 56 | 21 | 0 | 5 | 12 | 6 | 0 | 54 |

**Table S2**. Partial $\eta^2$ values for haldane numerators and their appropriate models for each study revisited. Darimont et al., (2009) not included because only darwins were used for analyses. Type refers to if the data were from a phenotypic study or a genotypic study (e.g., common garden). For *p*-values for question V., see Table S1.

| **Question** | **Model** | **N original** | **N updated** | **Explanatory variable** | **Effect size original** | **Effect size updated** | ***p-value* original** | ***p-value* updated** |
|---|---|---|---|---|---|---|---|---|
| I | Absolute Haldane numerators ~ generations * human influence | 2412 | 6233 | Generations | 0.006 | 0.001 | **< 0.001** | **0.031** |
| | | | | Human Influence | 0.072 | 0.0062 | **< 0.001** | **< 0.001** |
| III | Absolute Haldane numerators ~ introductions * generations | 104 | 320 | Generations | 0.196 | 0.302 | 0.858 | 0.744 |
| | | | | Introductions | 0.577 | 0.670 | 0.693 | 0.973 |
| V | Log absolute Haldane numerators ~ disturbance + type + years + generations | | 2015 | Generations | | 0.506 | | |
| | | | | Disturbance | | 0.892 | | |
| | | | | Type | | 0.0894 | | |

**Table S3.** Estimates, standard errors, Tukey test categorizations, and type II likelihood ratio test results for models predicting $\log_{10}$ absolute value haldanes. Chi-squared values are for the variable-specific likelihood ratio tests.

**$\log_{10}$ haldanes**

| | Variable | Est. | SE | Tukey | $\chi^2$ | df | *p* |
|---|---|---|---|---|---|---|---|
| Disturbance | Climate change | -0.84 | 0.07 | a | 144.83 | 6 | < **0.001** |
| | Hunting / harvesting | -0.46 | 0.06 | bc | | | |
| | Introduction | -0.47 | 0.05 | b | | | |
| | Landscape change | -0.45 | 0.11 | bc | | | |
| | Local response to introduced | -0.20 | 0.10 | cd | | | |
| | Natural | -0.50 | 0.06 | b | | | |
| | Pollution | 0.02 | 0.12 | d | | | |
| Phenotypic (vs. genetic) | | 0.08 | 0.05 | | 2.15 | 1 | 0.143 |
| $\log_{10}$ years | | 0.10 | 0.02 | | 21.91 | 1 | < **0.001** |
| $\log_{10}$ generations | | -0.19 | 0.04 | | 22.40 | 1 | < **0.001** |
